# Functional Divergence and Conservation in the QueC Protein Family (PF06508): From tRNA Modification to Anti-Phage Defense

**DOI:** 10.64898/2026.04.08.717055

**Authors:** Kaitlynn Libby, Rahul Benda, Valérie de Crécy-Lagard, Geoffrey Hutinet

**Author notes:** corresponding authors: GH -, VdC. KL: University of Florida, ICBR, Gainesville, FL 32610, RB: Evotec Biologics, Inc., Redmond, WA 98053.

## Abstract

The QueC protein family (PF06508) is best known for its role in the biosynthesis of 7⍰deazaguanine derivatives, including the tRNA modification queuosine and the DNA base 7⍰cyano-7⍰deazaguanine (preQ_0_). Recent discoveries, however, reveal that this ancient scaffold has been repeatedly repurposed for distinct biological functions, including roles in anti⍰phage defense. Here, we combine Sequence Similarity Networks (SSN) with genomic neighborhood analyses to map the functional diversification of the PF06508 superfamily. We delineate the canonical QueC cluster and experimentally define a refined catalytic signature, identifying previously unrecognized residues essential for tRNA modification. Beyond the canonical function, we characterize the evolutionary repurposing of the QueC fold in anti-phage systems, distinguishing between the “divergent specialist” QatC, which remodels the active site, and the “minimalist conservationist” Cap9, which retains the ancestral catalytic core. We further uncover the expansion of the superfamily into additional biochemical pathways, including cofactor biosynthesis (e.g., LarE) and purine salvage. Together, these findings provide a comprehensive framework for understanding how the QueC scaffold has been adapted from small⍰molecule biosynthesis to roles in protein modification and phage defense.

## Introduction

Derivatives of the 7-deazaguanine base are essential nucleic acid modifications ubiquitously distributed across the three domains of life and viruses [1]. These molecules serve two principal biological roles. First, they are critical components of transfer RNA (tRNA). In Bacteria and Eukarya, queuosine (Q) is a hypermodified nucleoside found at the wobble position (position 34) of tRNAs with GUN anticodons (Asp, Asn, His, Tyr). This modification is critical for translational fidelity and codon selection efficiency [1]. In Archaea, the analog archaeosine (G^+^) modifies position 14 or 15 in the D-loop of most tRNAs, stabilizing the tRNA’s tertiary structure [1]. Second, several 7-deazaguanine derivatives have been identified in the DNA of bacteria, bacteriophages, and archaeal viruses, where they function in defense against mobile genetic elements [2] or as anti-restriction systems [3, 4]. Despite these functional divergences, the biosynthetic pathways for both tRNA and DNA modifications share a common central intermediate: 7-cyano-7-deazaguanine (preQ_0_) [1].

In the well-characterized *de novo* bacterial pathway, preQ_0_ synthesis begins with GTP. This precursor is converted to 7-carboxy-7-deazaguanine (CDG) through a three-step enzymatic pathway [1]. This pathway is initiated by FolE (GTP cyclohydrolase I), an enzyme shared with the folate and biopterin biosynthesis pathways [5]. The product of FolE is subsequently converted to 6-carboxy-5,6,7,8-tetrahydropterin (CPH_4_) by QueD, which is the first dedicated step in the biosynthesis of natural 7-deazaguanine derivatives [6]. The radical S-adenosylmethionine (SAM) enzyme QueE catalyzes a complex rearrangement of CPH_4_ to form CDG [7]. The final step in preQ_0_ synthesis is catalyzed by QueC, a zinc-dependent metalloenzyme that converts CDG to preQ_0_ in two consecutive ATP-dependent reactions via a 7-amido-7-deazaguanine (ADG) intermediate, using ammonia as the nitrogen donor [8, 9]. Structurally, QueC belongs to the Rossman fold superfamily and has been characterized by two specific motifs: C(x)_8_CxxCxxC, which binds a zinc dication, and SGGxDS, which binds a phosphate group [9].

While QueC is defined by its role in tRNA and DNA modifications, recent discoveries have revealed that homologous proteins play critical roles in entirely different biological processes. Notably, QueC-like enzymes are core components of sophisticated anti-phage defense systems, including QatC in the QueC-like associated with ATPase and TatD DNase system (Qat) [10] or Cap9 in the type IV cyclic oligonucleotide-based anti-phage signaling system (CBASS) [11]. In these contexts, the QueC scaffold has been proposed to catalyze a protein modification reaction—covalent deazaguanylation of a partner protein—rather than to synthesize a base [12, 13]. In this work, we combine sequence similarity networks and genomic analysis to systematically map the functional landscape of the PF06508 superfamily, define the signature of canonical QueC, and explore the evolutionary trajectories of its functionally divergent paralogs.

## Material and Methods

### Strains and Plasmids generation

Primers for cloning and mutagenesis were designed using APE version 3.1.3 [14], synthesized by Eurofins (Louisville, KY), and referenced in **Supplemental Table 1**. pLG027 *qatC* [10] was cloned into the pBAD24 expression vector between the NcoI and SbfI restriction sites using Phusion High-Fidelity DNA polymerase, restriction enzymes, and T4 ligase from New England Biolabs (NEB, Ipswich, MA). Point mutations were generated using the Q5 Site-Directed Mutagenesis Kit from NEB. All plasmids were sequenced and maintained in *E. coli* DH5⍰, and assayed in *E. coli* MG1655 *queC* mutant from [3]. The QIAprep Spin Miniprep Kit from Qiagen (Hilden, Germany) was used to extract plasmids. All plasmids and strains are referenced in **Supplemental Table 2**, respectively. All *E. coli* were grown in LB. Plasmids were selected using 100 ug/mL of Ampicillin. Expression of the recombinant genes was repressed with Glucose 0.2% or induced with Arabinose 0.2%.

### QueC complementation Assay

QueC activity was measured using a Queuosine-modified tRNA gel shift assay as previously described [3]. Overnight bacterial cultures were diluted 1/100-fold into 5 mL of LB supplemented with 0.4% arabinose and 100 μg/mL ampicillin and grown for 2 h at 37 °C. Cells were harvested by centrifugation at 16,000 × g for 1 min at 4 °C. Cell pellets were immediately resuspended in 1 mL of Trizol (Life Technologies, Carlsbad, CA). Small RNAs were extracted using the PureLinkTM miRNA Isolation Kit from Invitrogen (Carlsbad, CA) according to the manufacturer’s protocol. Purified RNAs were eluted in 50 μL of RNase-free water, and tRNA concentrations were measured with a NanoDrop® ND-1000 Spectrophotometer (Thermo Fisher Scientific, Waltham, MA). RNA (500 ng) was migrated in a 10% acrylamide/ bisacrylamide (29:1) gel, containing Tris-EDTA acetate (TAE) 1X, Urea 8 M, and supplemented with 5 μg/mL 3-(acrylamido)-phenylboronic acid. RNA was transferred onto a BiodyneTM B Nylon membrane (0.45 μm, Thermo Scientific, Rockford, IL). tRNA samples were detected using a (5⍰biotin-CCCTCGGTGACAGGCAGG-3 ⍰) probe that anneals with tRNA_Asp_ ^(GUC)^ at a final concentration of 0.3 μM and the Chemiluminescent Nucleic Acid Detection Module Kit (Thermo Scientific, Rockford, IL), with the blocking buffer replaced by DIG Easy Hyp buffer (Roche, Mannheim, Germany).

### Sequence Similarity Network Generation

The Sequence Similarity Network (SSN) for the Pfam family PF06508 was generated in January 2023 using the Enzyme Function Initiative-Enzyme Similarity Tool (EFI-EST) web server [15, 16]. The “Families” option (Option B) was selected using the Pfam identifier PF06508 and the UniRef90 database to reduce redundancy. An alignment score of 30 was selected for the final network generation to strictly separate functionally distinct clusters. For visualization purposes, the network was filtered to a 65% sequence identity representative node set and visualized using Cytoscape version 3.10.3 [17]. Quantitative analysis of the network was performed on the full dataset (100% identity). Node attributes and metadata (**Supplemental Data 1**) were extracted using ssn2tsv (https://github.com/vdclab/ssn2tsv). Superkingdom distribution and sequence length analysis were performed manually on the extracted dataset using Microsoft Excel version 365.

### Multiple Sequence Alignments and Motif Analysis

Multiple sequence alignments for individual clusters were initially generated using the EFI-EST Cluster Analysis utility. To improve alignment quality and logo legibility, sequences were re-aligned using the MAFFT version 7 web server with default parameters [18, 19]. Sequence logos were generated using WebLogo 3 [20].

### Genomic Neighborhood and Functional Annotation

Genomic neighborhood networks were generated using the EFI Genome Neighborhood Tool (EFI-GNT)

[15] with an 80% sequence identity threshold. Detailed functional annotation of gene neighborhoods was performed manually by cross-referencing UniProt, Pfam, and InterPro databases. The PaperBLAST tool [21] was used to identify literature associated with specific sequences, and NCBI BLAST [22] was used to verify sequence identities. Additional analysis quantified the number of genes adjacent and in the same orientation as the gene of interest in each genome (“in operon”). All genome neighborhood information has been compiled in **Supplemental Data 2**.

### Structural Analysis

Conserved residues were mapped onto protein structures using UCSF ChimeraX version 1.11.1 [23]. Experimental structures were obtained from the RCSB Protein Data Bank [24] when available (e.g., *B. subtilis* QueC, PDB: 3BL5). For proteins lacking experimental structures, 3D models were generated using the AlphaFold3 online tool in the presence of ATP and Zn^2+^ (https://alphafoldserver.com). All structures from this study are available in the ChimeraX save file in **Supplemental Data 3**.

## Results and Discussion

### The PF06508 Superfamily is Dominated by Canonical QueC

To investigate the functional landscape of the PF06508 (QueC) family, a sequence similarity network (SSN) was generated using the EFI-EST (see methods, **Figure 1A**). At an alignment score threshold of 30, the superfamily segregated into numerous distinct clusters. The 10 largest clusters, which together accounted for 98.5% of all sequences, were selected for detailed analysis (**Figure 1B**).

**Figure 1.**
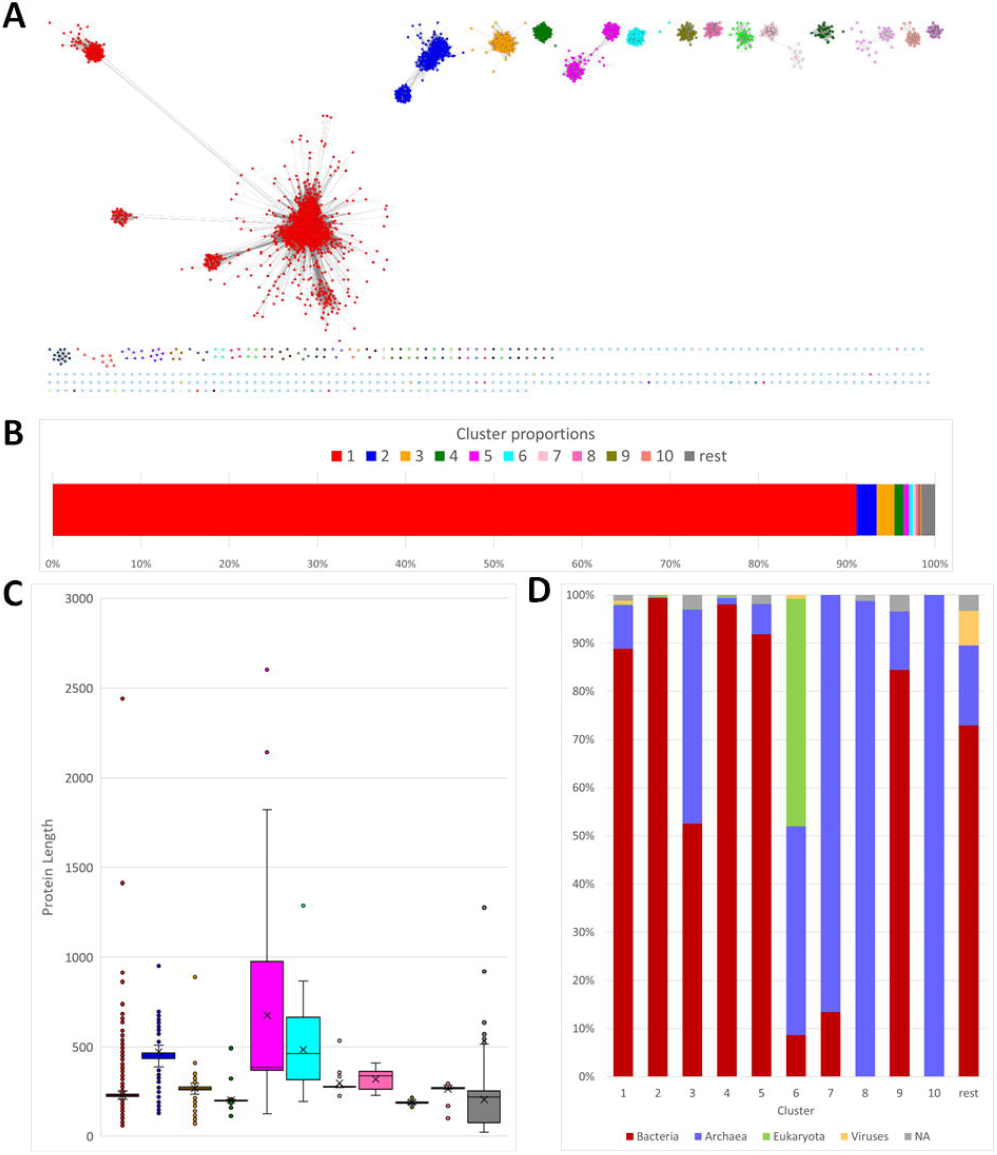
Non-homogeneity of PF06508 protein family. **(A)** Sequence Similarity Network (SSN) of the PF06508 family generated with an alignment score threshold of 30. Individual clusters are colored arbitrarily. **(B)** Distribution of sequences among the top 10 largest clusters, representing 98.5% of the dataset. **(C)** Protein length distribution across the major clusters. **(D)** Taxonomic distribution of the major clusters among Super-Kingdom and Viruses.

Cluster 1 is the dominant group, encompassing 91% of all sequences. This cluster’s median protein length of ∼222 residues (**Figure 1C**) is in excellent agreement with the size of experimentally characterized QueC orthologs, such as *Escherichia coli* QueC (231 aa) and *Bacillus subtilis* QueC (219 aa), which are both members of this cluster. In contrast, the other clusters show significant variation in median length (**Figure 1C**).

The taxonomic distribution (**Figure 1D**) further supports Cluster 1 as the canonical family. It is predominantly composed of sequences from Bacteria (∼89%) and Archaea (∼9%), the two domains known to possess the *de novo* 7-deazaguanine pathway. The family also contains a small number of Eukaryotic sequences, but these are almost exclusively found outside of Cluster 1 (e.g., in Cluster 6). The 43 Eukaryotic sequences found within Cluster 1 appear to be artifacts of horizontal gene transfer from bacterial symbionts or species misallocations from contaminants. For instance, the sequence from the ant *Lasius niger* shares 89.4% identity with a sequence from an *Acetobacteraceae bacterium*, a known ant gut symbiont (**Supplemental Table 3**). Similarly, the sequence from the kissing bug *Rhodnius prolixus* shows 85% identity to *Brenneria*, a bacterial pathogen (**Supplemental Table 3**). This indicates that authentic, functional QueC is absent in Eukarya, consistent with their queuosine auxotrophy.

Genomic neighborhood analysis provides further evidence (**Supplemental Table 4**). Members of Cluster 1 are frequently encoded adjacent to other genes in the pathway, such as *queD* (32.6 %, 30.9 % in operon) and *queE* (26.1 %, 21.7 % in operon). This conserved genetic linkage is a hallmark of the queuosine/archaeosine biosynthetic pathway and is not observed in the other major clusters. The combination of its dominant size, correct taxonomic profile, appropriate protein length, and conserved genomic context strongly identifies Cluster 1 as the *bona fide* QueC family.

### The Catalytic Signature of Canonical QueC

To analyze the spatial arrangement of catalytic residues, a structural model of the *E. coli* MG1655 QueC protein was generated using AlphaFold3 in the presence of ATP and zinc, both required for function [8, 9] (**Supplemental Figure S1A**). Building this model was required because the available experimental structure from *B. subtilis* (PDB: 3BL5) is partially cropped [9]. The *E. coli* model shows high structural homology and superimposes well with *B. subtilis* structure, validating its use as a reliable scaffold (**Supplemental Figure S1B**). This model was therefore used to map the conserved residues identified from a multiple-sequence alignment of all Cluster 1 sequences (**Figure 2A**) onto the structural model (**Figure 2BC and 3A**).

**Figure 2.**
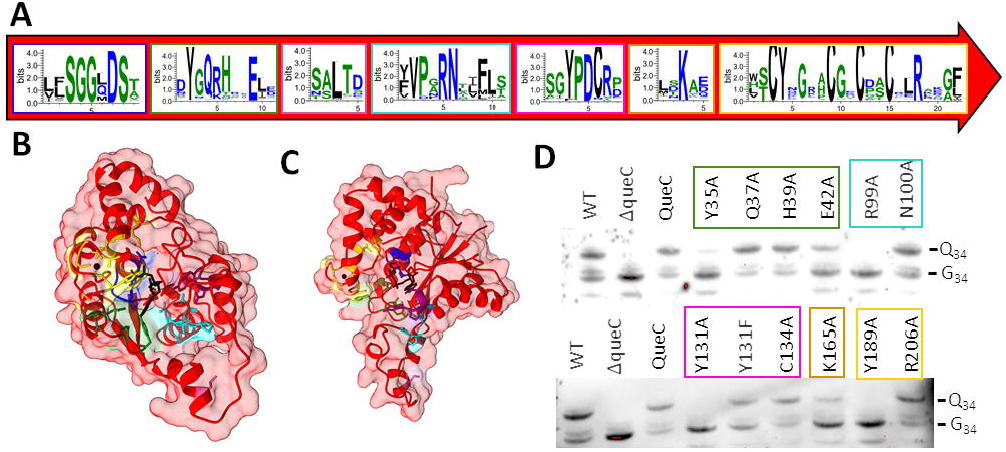
Catalytic signature of the canonical QueC. **(A)** Sequence logo of Cluster 1 derived from the multiple sequence alignment, highlighting conserved motifs ordered from N-terminal to C-terminal. The height of each letter indicates the degree of conservation. **(B and C)** AlphaFold3 model of *E. coli* QueC (UniProt ID: P0AG67) in complex with ATP and Zn^2+^ in two different orientations. Conserved residues identified in (A) are mapped onto the structure following the motif box coloring. **(D)** Northern blot analysis of tRNA extracted from an *E. coli* ΔqueC strain complemented with plasmids expressing wild-type (QueC) or mutant QueC variants (formatted as X###A), compared to the WT MG1655. The presence of the retarded band indicates successful formation of queuosine (Q_34_) in tRNA, compared to unmodified (G_34_).

Multiple sequence alignments confirmed the high conservation of the two hallmark motifs of the QueC family [9] at both extremities of the protein primary structure: the SGGxDS phosphate-binding loop (residues 11-16), consistent with QueC’s classification in the Rossman-fold superfamily, and the C(x)_8_CxxCxxC zinc-binding motif (residues 186-204). In addition to these known features, several other highly conserved motifs were identified: the YxQxHxxE motif (residues 35-42), which is located at the opening of the binding pocket and may interface with both the ATP and the 7-deazaguanine substrate; the VPxRN motif (residues 96-100), located close to the ATP in the model; and the YPDC motif (residues 131-134), at the opening of the binding pocket and likely to interact with the substrate. Two highly conserved single residues were also identified: L75, located in the pocket, and K165, on the surface of the protein.

To validate the functional importance of these newly identified conserved residues, site-directed mutagenesis of *E. coli* QueC and an *in vivo* complementation assay for the Q modification in *E. coli* tRNA were performed (**Figure 2D**). Mutagenesis of Y35A (in the YxQxHxxE motif) and R99A (in the VPxRN motif) resulted in a complete loss of function. Similarly, the Y131A mutant (in the YPDC motif) was inactive, although a Y131F mutant retained partial activity, suggesting a requirement for an aromatic ring at this position. These three residues orient toward the substrate pocket adjacent to the ATP (**Figure 3A**), and are essential for preQ_0_ formation from CDG. Furthermore, the mutagenesis data revealed that the Y189A mutant, located within the canonical zinc-binding region, is also completely inactive, highlighting a previously uncharacterized, essential role for this tyrosine in the catalytic mechanism.

**Figure 3.**
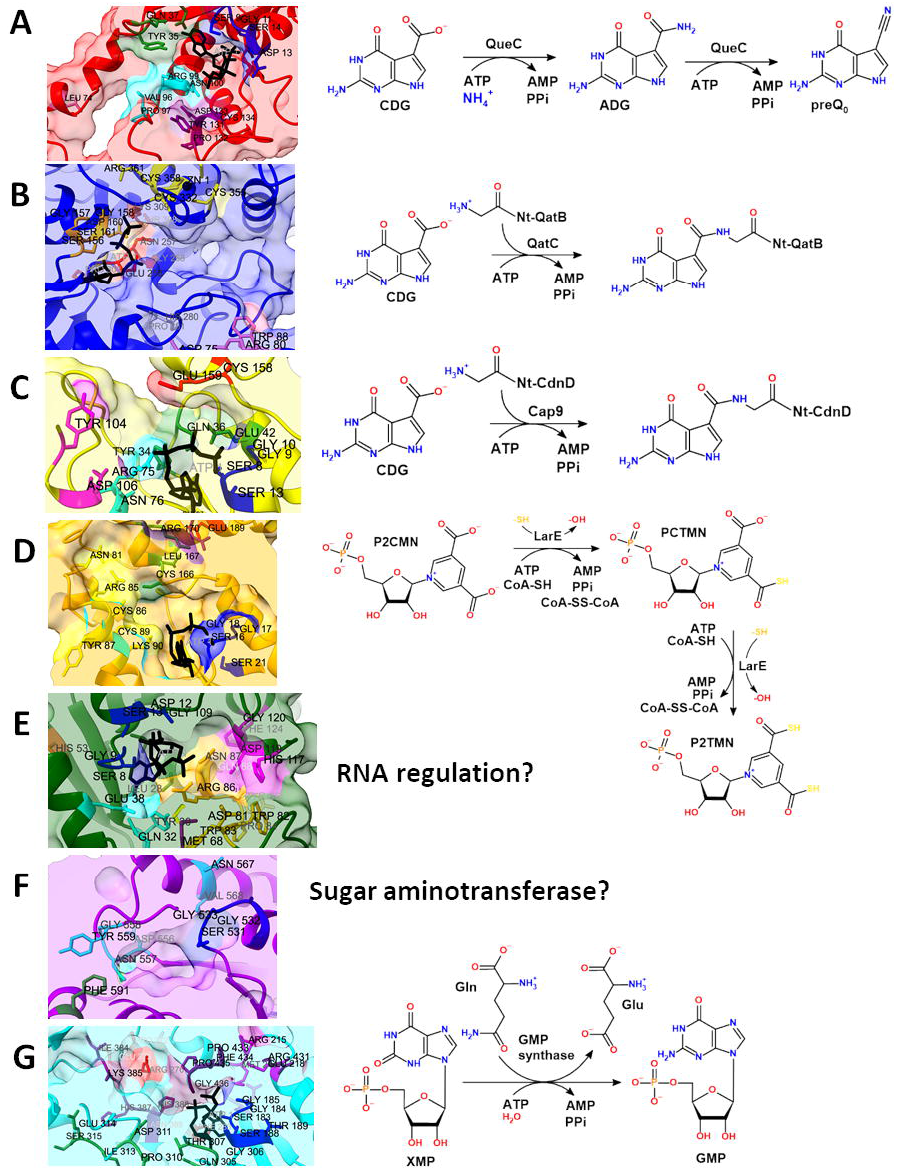
Proposed function of studied clusters. Catalytic core surrounding the SGGxDS motif and postulated catalytic reaction of Cluster 1 **(A)**, Cluster 2 **(B)**, Cluster 9 **(C)**, Cluster 3 **(D)**, Cluster 4 **(E)**, Cluster 5 **(E)**, and Cluster 6 **(G)**. Color of each structure by cluster color in **Figure 1**. Specific conserved residue colors are described in **Figure 2** for Cluster 1, and **Supplemental Figures 2** to **7** for Cluster 2, 9, 3, 4, 5, and 6 respectively. Abbreviations: *CDG*, 7-carboxy-7-deazaguanine; *ADG*, 7-amido-7-deazaguanine; *preQ*_0_, 7-cyano-7-deazaguanine; *ATP*, adenosine triphosphate; *AMP*, adenosine monophosphate; *PPi*, pyrophosphate; *Nt*, QatB or Cap9 N-terminal; *P2CMN*, pyridinium-3,5-biscarboxylic acid mononucleotide; *PCTMN*, pyridinium-3-carboxy-5-thiocarboxylic acid mononucleotide; *P2TMN*, pyridinium-3,5-bisthiocarboxylic acid mononucleotide; *CoA*, Coenzyme A; *XMP*, xanthosine monophosphate; *GMP*, guanosine monophosphate; *Gln*, L-glutamine; *Glu*, L-glutamate.

### Repurposing of the QueC Scaffold for Anti-Phage Defense

The analysis of paralogous clusters revealed that the QueC scaffold has been independently repurposed for roles in anti-phage defense on at least two occasions. This evolutionary co-option pivots the enzyme’s chemistry from the synthesis of a small molecule to the catalysis of a novel protein modification.

Cluster 2 contains the QatC protein, a core component of the Qat defense system [10], and is mainly associated with Bacteria (**Figure 1D**). Genomic neighborhood analysis (**Supplemental Table 4**) reveals that Cluster 2 genes are clustered with *qatA* (ATPase, PF07693, 34.3%, 33.8% in operon) and *qatD* (TatD-like DNase, PF01026, 33.6%, 32.4% in operon). Although the gene encoding the system’s substrate protein, *qatB* [13], is not covered by an existing Pfam model, it is found in the operon at a similar frequency to the other genes (data not shown). Structurally, the core of QatC preserves the canonical QueC Rossman-fold but is elaborated with unique N- and C-terminal extensions that account for the increased median length (∼400 aa, **Supplemental Figure 2**). Examination of the conserved residues reveals a striking pattern of conservation and divergence compared to Cluster 1. While the ATP-binding SGGxDS loop is strictly conserved, the specific catalytic residues defined for Cluster 1—including YxQxHxxE, VPxRN, and YPDC—are notably absent. Instead, Cluster 2 (residues numbering from the *E. coli* sequence used in [10], **Supplemental Figure 2**) is defined by a unique set of conserved motifs: D(x)_4_R(x)_7_WxR (residues 75 to 90), DxW/YxxxF (residues 121 to 127), RxRxxxF/Y (residues 209 to 214), ExG (E252, G258), HP (H280, P281), P(x)_4_TK (residues 303 to 309), and conserved single residues D63, previously identified as a catalytic residue [13], and Y376. The zinc-binding motif is also retained but presents as a variant C(x)_19_CGxCxxRR pattern (residues 332 to 362). This active site remodeling reflects the enzyme’s specialized function: instead of binding a small molecule, the repurposed scaffold has been proposed to accommodate the glycine N-terminus of QatB to catalyze its ATP-dependent protein carboxydeazaguanylation (**Figure 3B**), an essential activation step for the anti-phage response [13]. Consistent with this shift in substrate specificity, we found that expression of QatC failed to restore the Q-modification in an *E. coli* Δ*queC* strain (data not shown), confirming that it has lost the ability to synthesize preQ_0_.

Cluster 9 contains the Cap9 proteins, the signature effector of Type IV CBASS systems [11], which function as ATP-dependent ligases catalyzing the N-terminal carboxydeazaguanylation of their CD-NTase partner proteins. This modification is the critical switch that activates the synthesis of cyclic oligonucleotide second messengers, thereby initiating the immune response. Genomic neighborhood analysis (**Supplemental Table 4**) reveals that Cluster 9 proteins are encoded adjacent to the gene for the system’s sensor, a nucleotidyltransferase (CD-NTase, PF18144, 67%, 65% in operon). Structurally, Cap9 represents the minimal catalytic unit of the superfamily, with a median length of only ∼185 residues (**Figure 1C**). Unlike the extended Cluster2/QatC scaffold, Cluster 9 appears to be a truncated Rossman-fold. However, sequence analysis reveals a remarkable preservation of the canonical active site architecture (**Supplemental Figure 3**). Unlike Cluster 2 (QatC), which has completely remodeled its catalytic center, Cluster 9 retains the core catalytic signature of the canonical QueC (Cluster 1). In addition to the defining motifs, LxSGGxDS (residues 6 to 13) and CE(x)_6_CxxCxxCxD (residues 158 to 174), the signature YxQxHxxE motif is conserved as a Y/FxQ(x)_5_E variant (residues 34 to 42), as are the YPDC substrate-binding motif is preserved as H(x)_4_YxDCxxxF (residues 99 to 111), and the RN motif (residues 75 and 76), part of the VPxRN loop, and PF(x)_4_K (residues 132 to 138), likely analogous to the conserved K of Cluster 1. This suggests that Cap9 and QatC represent two distinct evolutionary strategies for achieving the same biological goal (protein deazaguanylation). QatC (Cluster 2) is a “divergent specialist” that evolved novel motifs to accommodate its function, whereas Cap9 (Cluster 9) is a “minimalist conservationist,” stripping away non-essential structural elements while strictly maintaining the ancestral catalytic machinery. Thus, while Cluster 9 shares its biological function with Cluster 2 (**Figure 3C**), its active site topology closely resembles the canonical metabolic enzyme, QueC (Cluster 1).

### Functional Diversification into Other Metabolic Pathways

Beyond the canonical QueC and its anti-phage paralogs, the remaining clusters illustrate how the superfamily’s catalytic core has been adapted for diverse metabolic functions. In these lineages, the conserved ATP-binding machinery is generally retained, but the substrate-recognition motifs and metal-binding sites have diverged significantly to accommodate distinct chemical reactions.

Cluster 3 corresponds to the LarE family, involved in the synthesis of the cofactor for lactate racemase [25]. Genomic neighborhood analysis (**Supplemental Table 4**) strongly supports this assignment: Cluster 3 sequences are encoded in bacteria and archaea (∼53% and ∼44% respectively, **Figure 1D**) adjacent to LarC (Ni_insertion, PF01969, ∼54%, ∼52% in operon) and LarB (AIRC, PF00731, ∼47%, ∼29% in operon).

While the SGGxDS ATP-binding motif is preserved, the C(x)_8_CxxCxxC zinc-binding motif of the canonical QueC is absent (**Supplemental Figure 4**). Instead, Cluster 3 is defined by a strictly conserved three-cysteine motif (**Figure 3D**). Recent structural and biochemical characterization of non-sacrificial LarE revealed that these three cysteines coordinate a [4Fe-4S] cluster binding to a thiol group used for the reaction, whereas the sacrificial LarE possesses only one conserved cysteine that is consumed into a serine [25, 26]. This marks a fundamental mechanistic shift from the zinc-dependent ligation of QueC to [4Fe-4S]-dependent sulfurtransferase activity. Furthermore, none of the canonical QueC motifs identified in this study are present. Structurally, Cluster 3 aligns poorly with QueC, exhibiting significant divergence outside the core Rossmann fold (**Supplemental Figure 4**).

Cluster 4 was found primarily within *Proteobacteria* (>2/3 of sequences). While the protein sequence is slightly shorter than the canonical QueC by approximately 10 residues, the N-terminal region retains significant conservation. It shares the SGGxDS ATP-binding loop and a slightly different YGQ(x)_6_E motif. Structural modeling suggests the N-terminus aligns well with the QueC fold (**Supplemental Figure 4**). However, the rest of the protein is defined by a unique set of conserved motifs distinct from the canonical enzyme, and a variant of the C-terminal cysteine-rich signature (CH(x)_5_CxxCxGCxK). This implies a retained ATP-dependent mechanism but a divergent substrate specificity (**Figure 3E**). Nearly every neighborhood (**Supplemental Table 4**) contains a PfkB domain-containing protein (PF00294-PF05014, ∼89%, ∼88% in operon), likely a nucleoside 2-deoxyribosyltransferase involved in purine salvage. Cluster 4 proteins frequently co-occurred with Helix-Turn-Helix domains, suggesting a potential role in regulating expression, possibly at the RNA level [27, 28].

Cluster 5 segregates into two distinct subclusters (**Figure 1A**) characterized by significant functional heterogeneity, despite sharing a conserved residue signature comprising a modified P-loop SGGxDSxY, a cysteine-rich CxxC(x)_6_CxC motif, and a potential substrate-binding YDC triad, although they appear in reverse sequential order compared to canonical QueC (**Supplemental Figure 6**). Subcluster 1 is composed of massive multi-domain proteins (ranging from 900 to 2603 amino acids) predominantly found in *Flavobacteria*. In contrast, Subcluster 2 consists of smaller proteins (300-400 amino acids) annotated as N-sugar amidotransferases (**Figure 3G**). A key representative is WbpG, a gene essential for the synthesis of the O-antigen lipopolysaccharide (LPS) in *Pseudomonas aeruginosa* [29].

Cluster 6 is firmly identified as the GMP Synthase (GMPS) family. In sharp contrast to the bacterial QueC, this cluster is dominated by sequences from Eukaryota and Archaea (**Figure 1D**). Structural diversity within this cluster correlates with taxonomy: Eukaryotic members are typically large (>630 aa), reflecting the fusion of the QueC-like ATP-pyrophosphatase domain with a glutamine amidotransferase (GATase) domain. Conversely, Archaeal members, particularly from *Thermoplasmata*, are shorter (∼315 aa) and appear to function as a standalone subunit, with the GATase subunit encoded separately downstream in the genome neighborhood (PF00117, ∼26%, ∼22% in operon, **Supplemental Table 4**). Conserved residue analysis (**Supplemental Figure 7**) reveals a distinct signature for this lineage. The ATP-binding P-loop is preserved as SGGVDSTV. However, the canonical QueC motifs are replaced by Cluster 6-specific sequences including DxGxM/LRxxE, QGTxxPDxIES, IKxHHN, EPL, KDEVR, and RxPFPG. These motifs likely facilitate the specific binding of Xanthosine Monophosphate (XMP) and the coupling of ATP hydrolysis with ammonia transfer (**Figure 3G**), a mechanism distinct from the preQ_0_ synthesis of QueC. Notably the cysteine-rich zinc binding motif is absent in this cluster.

Finally, Clusters 7, 8, and 10 represent lineages that are almost exclusively restricted to Archaea, with a notably high prevalence in methanogens. Given the relatively small size of those clusters, we did not extend our detailed sequence or neighborhood analysis to these clusters. They likely represent further diversifications of the QueC-like fold for archaea-specific functions, but their precise roles remain to be determined.

## Conclusions

Our comprehensive mapping of the PF06508 superfamily illustrates the remarkable evolutionary plasticity of the QueC Rossmann-fold. We successfully established the catalytic signature of canonical QueC and identified previously overlooked residues essential for tRNA modification. Beyond primary metabolism, our analysis highlights how this versatile ATP-dependent scaffold has been repeatedly co-opted to drive biological innovation. From the structural remodeling required for anti-phage protein deazaguanylation to the substitution of zinc coordination for sacrificial sulfur transfer, these divergent clusters exemplify nature’s ability to repurpose existing enzymatic architecture. Resolving the precise biochemical roles of the remaining uncharacterized clusters promises to reveal further novel chemistries. Ultimately, this framework not only clarifies the functional boundaries of the QueC family but also provides a robust, predictive foundation for future experimental characterization of diverse metabolic and immune pathways.

## Supplemental data

Supplemental figures, tables, and data have been deposited in figshare (https://figshare.com/s/694e3add2ef9f19a906c).

## Acknowledgements and disclaimer

This work was funded by the National Institutes of Health (Grants R01GM70641 and R35GM156215 to V dC-L). The content is solely the responsibility of the authors and does not necessarily represent the official views of the National Institutes of Health. Generative AI tools were utilized solely for the purpose of copy-editing and refining the text of this manuscript. The authors confirm that AI was not used to generate, process, or analyze any data presented in this study beyond what is described in the methods.

